# DCAFA: Differential Community Abundance and Feature Analysis for Histological Images

**DOI:** 10.64898/2026.04.28.721329

**Authors:** George Wright, Piotr Keller, Joanne Muter, Jan Brosens, Sabine Tejpar, Fayyaz Minhas

**Affiliations:** Predictive Systems in Biomedicine (PRISM) Lab, Department of Computer Science, University of Warwick, Coventry, UK; Warwick Medical School, Division of Biomedical Sciences, University of Warwick, Coventry, UK; Laboratory for Digestive Oncology, KU Leuven, Leuven, Belgium

**Keywords:** Regression-based Modelling, Community Abundance Analysis, Computational Pathology, Phenotyping

## Abstract

Histological images are composed of diverse local structures such as cells, glands, or tissue patches, whose organisation and relative abundance reflect underlying biological processes. While most computational approaches focus on analysing individual features, many clinically relevant signals arise from changes in the composition of these structures rather than isolated measurements. This motivates the need for explicitly modelling how groups of similar instances vary across samples and relate to outcomes. We present DCAFA (Differential Community Abundance and Feature Attribution Analysis), a regression-based framework for analysing hierarchical biomedical data through both compositional and feature-level perspectives. DCAFA groups instances into latent communities representing recurring morphological or phenotypic patterns, and then performs two complementary analyses: (i) community composition analysis, which identifies groups that are enriched or depleted across outcomes, and (ii) feature attribution analysis, which quantifies how instance-level features relate to outcomes directly or within specific communities. Both use generalised linear and mixed-effects models, enabling covariate adjustment and inference through effect sizes, confidence intervals, and false discovery rate control. We demonstrate the utility of DCAFA across multiple biomedical settings, including endometrial histopathology, spatial transcriptomics, multiplex immunofluorescence imaging, and predefined cell-type analyses in colorectal cancer. These examples identify interpretable compositional shifts and context-specific feature associations that are not captured by conventional feature-based approaches. By unifying differential abundance testing and feature attribution within a single statistical framework, DCAFA serves as an openly available toolbox that provides a practical and interpretable means of linking tissue composition with clinical or molecular outcomes in biomedical imaging data.

Code available at: https://github.com/wgrgwrght/DCAFA

## 1 Introduction

Biological tissue samples are composed of diverse cellular and structural elements whose organisation reflects both normal physiology and disease. In histological images and related high-dimensional biomedical data, these elements do not occur in isolation but form groups of similar instances, such as morphologically or functionally related cells, glands, or tissue regions/patches. The relative abundance and composition of such groups encode key biological processes and are often indicative of pathological state [16]. Understanding how the composition of these groups varies within and across samples is therefore central to interpreting tissue structure and function. With the advent of high-throughput imaging and spatial profiling technologies, including histopathology, multiplexed imaging, and spatial or single-cell transcriptomics, it is now possible to characterise tissue organisation at unprecedented resolution [3]. However, translating these high-dimensional representations into interpretable and quantitative insight remains a fundamental challenge: *given a histology sample, how can we systematically characterise its composition and relate it to biological outcomes or conditions?*

Most existing analytical approaches do not explicitly model composition. Instead, they operate at the level of individual features, such as pixel intensities, molecular measurements, or learned embeddings, without directly capturing how instances assemble into higher-order structures [14]. As a result, biologically meaningful patterns must be inferred indirectly, often relying on complex models or heuristics. In contrast, many pathological processes manifest as shifts in the prevalence of tissue components, for example, changes in immune infiltration, glandular organisation, or stromal content, which may not be detectable through feature-wise analyses alone [5]. This motivates the need for methods that treat *composition as a primary object of analysis*, explicitly modelling how groups of similar instances change across conditions.

Differential abundance (DA) methods provide a statistical framework for identifying such compositional changes and have been widely used in single-cell and spatial omics. Approaches such as MILO [2], DA-seq [15], and edgeR-based models [9] detect groups of cells or neighbourhoods whose abundance differs across conditions. However, these methods are primarily designed for count-based inference within predefined or graph-derived groups and have seen limited adaptation to computational pathology. More importantly, they focus exclusively on detecting enriched or depleted populations and do not model how instance-level features relate to either group structure or outcomes. This limitation is particularly relevant in histopathology and related domains, where data are inherently hierarchical, with instances (e.g., cells or patches) nested within images or patients, and both the composition of groups and the properties of individual instances contribute to clinically meaningful signals.

To address this, we introduce DCAFA (Differential Community Abundance and Feature Attribution Analysis), a unified statistical framework for analysing hierarchical biomedical data through instance grouping. Here, we use the term *community* to denote groups of similar instances defined in feature space, which may optionally reflect spatial structure but are not restricted to it. DCAFA or-ganises instances into such communities based on morphological, molecular, or learned representations, and jointly addresses two complementary questions: (i) which communities are differentially abundant across conditions, and (ii) which instance-level features explain both community membership and outcome associations. By embedding these analyses within a generalised linear modelling framework, DCAFA enables statistically rigorous inference on both compositional shifts and feature-level effects, while naturally accommodating both bag-level outcomes (e.g., patient labels) and instance-level outcomes (e.g., cell states). To facilitate adoption in practice, we additionally provide an easy-to-use Python framework that enables researchers to run DCAFA directly on their own data and communities, supporting reproducible and accessible analysis across a wide range of biomedical applications.

The main contributions of this work are as follows:

1. We introduce DCAFA, a unified regression-based framework for analysing hierarchical biomedical data through community abundance and feature association, allowing both compositional analysis and feature-level inference within a single statistical formulation.
2. We formulate four complementary analysis settings within the same framework, covering bag-level and instance-level community abundance modelling as well as bag-level and instance-level feature association, thereby enabling inference across multiple levels of biological organisation.
3. Support flexible community definitions, including predefined categories, hard clustering, soft memberships, and overlapping neighbourhood-based groupings, making the framework applicable to a wide range of data.
4. We provide statistically principled modelling of community abundances and feature effects using generalised linear and mixed-effects models, with support for diverse response types, covariate adjustment, within-bag dependence, multiple testing correction, and compositional constraints such as offsets and collinearity handling.
5. DCAFA yields biologically interpretable insights across multiple biomedical applications, including endometrial histopathology, colorectal cancer spatial transcriptomics, multiplex immunofluorescence imaging, and predefined cell-type abundance analysis in colorectal cancer treatment response.

## 2 Methods

### 2.1 Problem Formulation

Histological images consist of heterogeneous mixtures of local structures, such as cells, glands, or tissue patches, whose relative abundance and characteristics vary across samples. These structures can be extracted using computational pipelines, for example via segmentation models, object detection, or systematic tiling of whole-slide images. Each extracted element is represented by a feature vector capturing morphological, spatial, or learned properties.

We represent each image as a collection of such elements and formalise this setting using a bag-instance framework. Specifically, each image *i* ∈{1, …, *B*} is treated as a *bag* containing multiple instances. Let *x*_*p*_ ∈ ℝ^*d*^ denote the feature vector of instance *p*, and let *s*(*p*) map each instance to its parent image. This induces a hierarchical structure in which instances are nested within images.

Outcomes or condition targets may be defined at different levels:

- image-level outcomes *y*_*i*_ (e.g., diagnosis, risk score, or gene expression),
- instance-level outcomes *t*_*p*_ (e.g., marker intensity or cell state).

In addition, each image may be associated with covariates *v*_*i*_ capturing technical or biological variation such as batch effects, acquisition site, patient group, or other control variables.

A central challenge is to relate these outcomes to the organisation of instances within each image. Direct modelling at the level of individual features is often insufficient, as biologically meaningful variation frequently arises from shifts in the *composition* of structural elements rather than changes in isolated measurements. For example, disease progression may manifest as changes in the prevalence of specific cell types or morphological patterns, even when individual feature distributions remain similar.

To capture these compositional effects, we propose **DCAFA** (Differential Community Abundance and Feature Attribution Analysis), a regression-based framework that introduces an intermediate representation based on groups of similar instances. Given instance-level feature representations, we first group instances into latent *communities* defined in feature space. These communities capture recurring visual or biological patterns shared across images and provide an interpretable representation of tissue organisation.

Within this representation, DCAFA addresses two complementary questions:

1. **Community composition (abundance):** which groups of similar instances are differentially abundant across outcomes or conditions?
2. **Feature attribution:** which instance-level features explain outcome variation, either directly or through their association with specific communities?

These analyses can be performed at either the instance or image level, yielding four related model families within a unified framework. We adopt a regression-based formulation, providing a principled and flexible way to model relationships between outcomes, community composition, and instance-level features while adjusting for covariates and hierarchical dependencies. This approach enables direct estimation of effect sizes via regression coefficients and statistically grounded inference through hypothesis testing and confidence intervals, with within-image dependencies handled by hierarchical modelling. Statistical significance is assessed using multiple testing correction to control the false discovery rate.

### 2.2 Community Construction and Representation

Given instance-level feature representations, our goal is to capture recurring patterns that reflect shared morphological or phenotypic characteristics across images. To this end, we group instances into *K communities* in feature space, each representing similar cells, glands, or patches.

Communities can be obtained through an unsupervised grouping of instance features 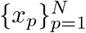. The framework is intentionally agnostic to the specific method used, and can accommodate a range of approaches including centroid-based clustering (e.g., *k*-means), probabilistic models (e.g., Gaussian mixture models), graph-based methods, or clustering applied to learned representations from deep neural networks. The choice of method, as well as the number and granularity of communities, is left to the user and may be adapted to the data modality and biological question of interest.

We denote by *m*_*pk*_ the membership of instance *p* in community *k*. The formulation supports both hard (*m*_*pk*_ ∈ {0, 1}) and soft memberships (*m*_*pk*_ ∈ [0, 1]), enabling communities to represent either discrete or overlapping patterns.

For each image *i*, community composition is summarised by aggregating instance memberships:

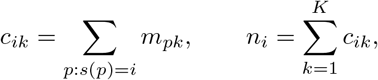

where *c*_*ik*_ denotes the abundance of community *k* in image *i*, and *n*_*i*_ is the total number of instances extracted from that image. This representation provides a compact and interpretable summary of tissue composition, capturing how different structural patterns are distributed within each sample.

Under hard assignments, the constraint 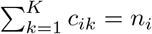 induces perfect collinear-ity between community abundances. To ensure identifiability in downstream regression models, one community is omitted as a reference category, and all estimated effects are interpreted relative to this baseline. In the case of soft assignments, approximate collinearity may still arise and is handled within the regression framework.

### 2.3 Bag-Level Community Composition Analysis

In many histopathology settings, labels are available only at the image or patient level, while the underlying signal is driven by the composition of structures within the image. For example, an image-level outcome *y*_*i*_ (e.g., diagnosis, risk score, or gene expression) may depend on the relative prevalence of specific morphological or cellular patterns.

To quantify these relationships, we model community abundances using a negative binomial regression. This choice reflects the discrete and overdispersed nature of instance-derived counts, while allowing principled modelling of relative composition. For each image *i* and community *k*, we define:

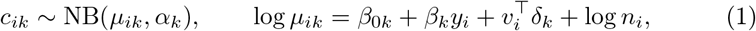

where *c*_*ik*_ denotes the abundance of community *k* in image *i, v*_*i*_ represents image-level covariates, and *n*_*i*_ is the total number of instances extracted from that image. The offset term log *n*_*i*_ ensures that the model captures *relative* rather than absolute abundance, thereby accounting for variation in the number of detected instances across images.

The coefficient *β*_*k*_ quantifies how the outcome *y*_*i*_ is associated with changes in the relative abundance of community *k*. Specifically, exp(*β*_*k*_) represents the multiplicative change in expected abundance for a one-unit increase in *y*_*i*_, conditional on covariates. For example, if *y*_*i*_ is a binary indicator of disease status and exp(*β*_*k*_) = 1.5, this indicates that community *k* is expected to be 50% more abundant (relative to total instances) in diseased samples compared to controls. Conversely, exp(*β*_*k*_) *<* 1 indicates depletion.

The dispersion parameter *α*_*k*_ accounts for variability exceeding that of a Poisson model, which commonly arises due to heterogeneity in instance detection and biological variation.

#### Algorithm 1

DCAFA Pipeline

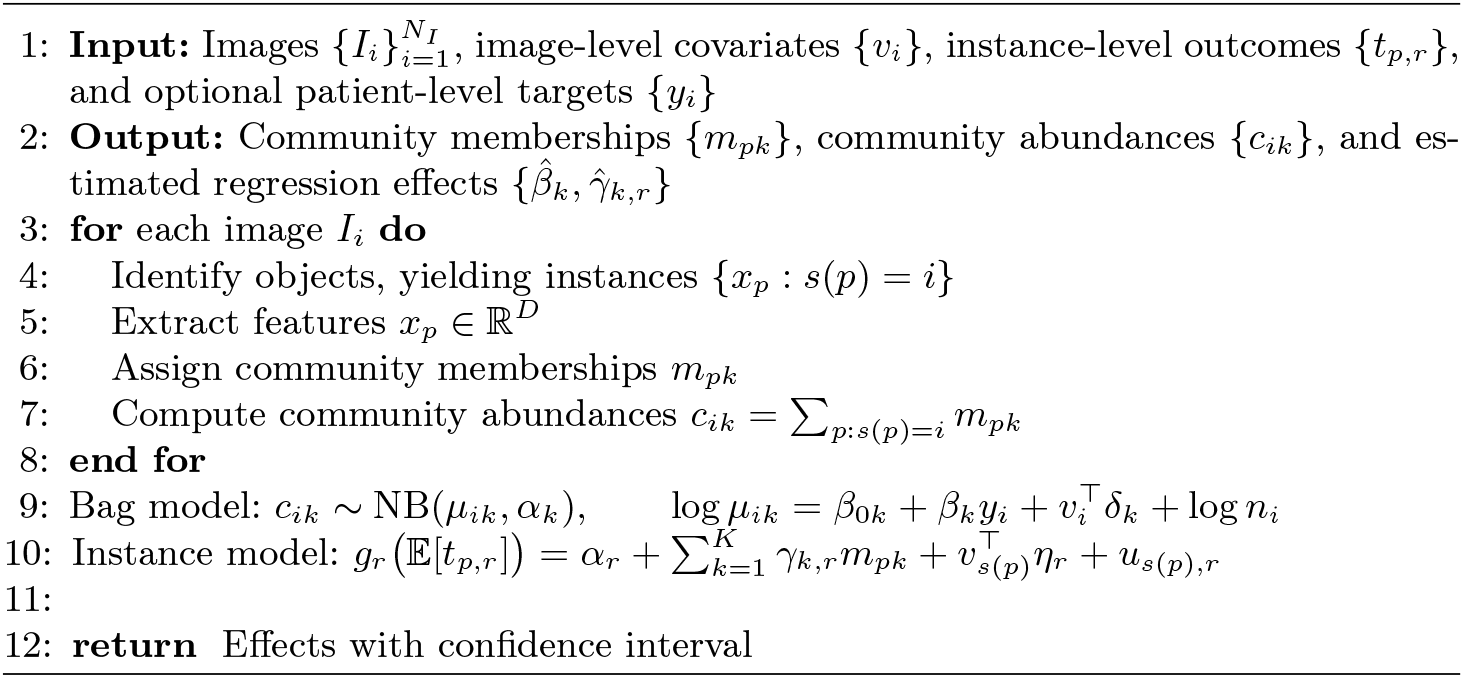

Positive values of 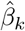 indicate enrichment of community *k* with increasing *y*_*i*_, while negative values indicate depletion relative to the reference community. This formulation is related to differential abundance approaches such as MILO [2], which also model neighbourhood-level counts using negative binomial regression. However, here communities are defined in feature space and integrated within a broader regression framework that enables joint modelling of composition and feature-level effects.

### 2.4 Instance-Level Community Composition Analysis

In some settings, outcomes are defined at the level of individual instances rather than entire images. For example, one may wish to relate cell-level measurements (e.g, marker intensities, morphological scores, or pathway activation indicators) to underlying structural patterns captured by communities. In such cases, the objective is to assess whether membership in different communities is associated with systematic variation in instance-level outcomes.

To quantify these relationships, we model instance-level responses using generalised linear mixed models. For each instance *p* and outcome component *r*, we define:

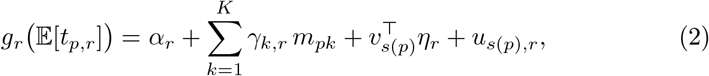

where *t*_*p,r*_ denotes the response associated with instance *p*, and *m*_*pk*_ encodes its membership in community *k*. Covariates *v*_*s*(*p*)_ capture image-level factors such as batch effects or patient metadata, and 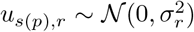. is a random intercept accounting for correlations among instances originating from the same image.

The link function *g*_*r*_(·) is selected according to the outcome type, for example identity for continuous measurements, logit for binary outcomes, or log for count-based responses.

The coefficients *γ*_*k,r*_ quantify how membership in community *k* is associated with changes in the expected value of the outcome. Intuitively, these coefficients describe how different groups of similar instances differ in their functional or morphological properties. For example, if *t*_*p,r*_ represents a protein expression level and *γ*_*k,r*_ is positive, this indicates that instances belonging to community *k* tend to exhibit higher expression of that protein relative to the reference community, after adjusting for covariates. Conversely, negative coefficients indicate reduced expression or prevalence.

By modelling community membership and instance-level outcomes jointly, this formulation enables direct interpretation of communities in terms of their biological or morphological characteristics, linking abstract groupings in feature space to measurable functional properties.

Complementary to community-based analysis, we directly assess how instance-level features relate to outcomes. This enables identification of the specific morphological, molecular, or learned characteristics that drive variation in instance-level responses.

We model instance-level outcomes using a generalised linear mixed model:

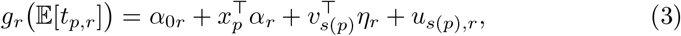

where *x*_*p*_ ∈ ℝ^*d*^ is the feature vector of instance *p, v*_*s*(*p*)_ are image-level covariates, and *u*_*s*(*p*),*r*_ is a random intercept accounting for correlations among instances from the same image. The link function *g*_*r*_(·) is chosen according to the outcome type.

The coefficients *α*_*r*_ quantify feature–outcome associations. In particular, *α*_*j,r*_ represents the change in the expected outcome associated with a unit increase in feature *j*, conditional on covariates. For example, if *t*_*p,r*_ represents marker expression, a positive coefficient indicates that higher values of the corresponding feature (e.g., nuclear size or staining intensity) are associated with increased expression. For non-Gaussian outcomes, exponentiated coefficients correspond to multiplicative effects.

Statistical significance is assessed using hypothesis tests on the regression coefficients, with multiple testing correction applied across features to control the false discovery rate. This formulation provides a direct and interpretable mapping from instance-level features to outcomes, complementing community-based analysis by identifying the attributes underlying observed compositional effects.

### 2.5 Bag-Level Feature Attribution Analysis

To relate instance-level features to image-level outcomes, local information must be propagated from instances to their parent images. We consider three complementary strategies that differ in how feature information is summarised across each image.

#### Instance-level modelling with shared labels

Under a weak supervision setting, the image-level outcome is assigned to all instances from the same image (*t*_*p*_:= *y*_*s*(*p*)_.) after which we then fit a mixed-effects model

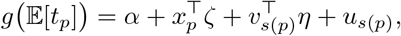

where *v*_*s*(*p*)_ are image-level covariates and *u*_*s*(*p*)_ is an image-specific random intercept. This random effect is necessary because all instances from the same image share the same label and are therefore not independent. The coefficients *ζ* quantify how instance-level features are associated with the image-level outcome, while accounting for within-image dependence.

#### Global aggregation

Alternatively, instance-level features can be averaged within each image:

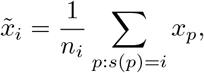

and used in a regression model

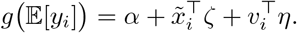

Here, *ζ* quantifies how the overall feature composition of an image relates to the outcome. This approach is simple and interpretable, but may obscure heterogeneity within the image.

#### Community-aware aggregation

To preserve intra-image heterogeneity, features can instead be aggregated within communities:

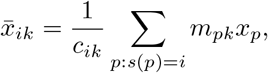

and modelled as

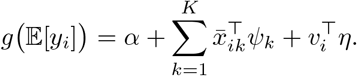

Here, *ψ*_*k*_ is a community-specific coefficient vector, so that the same feature may have different associations with the outcome in different communities. This enables detection of context-specific effects that may be masked by global averaging. Communities with *c*_*ik*_ = 0 are treated as missing for the corresponding summaries.

Across all three strategies, regression coefficients quantify how image-derived features are associated with image-level outcomes, either globally or within specific communities. For example, a positive coefficient indicates that larger values of a feature are associated with higher expected outcome values on the scale of the chosen link function.

### 2.6 Statistical Inference and Visualization

All models are estimated within a regression framework, enabling direct quantification of effect sizes and statistical significance. For each parameter of interest, inference is performed using Wald tests on the corresponding regression coefficients. Resulting *p*-values are adjusted for multiple comparisons using the Benjamini–Hochberg procedure to control the false discovery rate, ensuring robust identification of significant associations across communities and features.

Effect sizes are reported as regression coefficients and interpreted on the scale of the chosen link function. For non-Gaussian models, coefficients are exponentiated to yield interpretable measures such as odds ratios (for binary outcomes) or rate ratios (for count data), facilitating direct comparison across analyses. Confidence intervals are computed to characterise uncertainty in the estimated effects.

To support interpretation, results are visualised using complementary approaches. Forest plots are used where feasible to display effect sizes alongside confidence intervals, providing a clear view of both magnitude and uncertainty for individual associations. When the number of parameters is large, heatmaps are employed to summarise patterns of association across communities, features, and outcomes. These heatmaps are typically normalised to highlight relative differences and facilitate comparison across groups.

Together, these visualisation strategies provide a coherent and interpretable view of both community composition and feature-level effects. By combining statistically grounded inference with structured visual summaries, the framework enables identification and interpretation of patterns across multiple levels of biological organisation, linking instance-level characteristics, community structure, and sample-level outcomes.

## 3 Experiments and Results

### 3.1 Compositional Shifts in Endometrial Tissue

We first set out to examine glandular and nuclear changes in endometrial tissue using DCAFA. The human endometrium is known to undergo substantial remodelling following ovulation, reflected by coordinated structural and molecular changes linked to gene expression [10]. We analysed 624 whole-slide images (WSIs) stained with haematoxylin and CD56 immunohistochemistry, paired with log-normalised RT-qPCR gene expression data. Using ASTER [13], we segmented 233,113,551 nuclei and 529,569 gland components across the dataset. For downstream processing, 1,000 cells per WSI were sampled to balance computational efficiency with representation of intra-slide variability. From each segmented object, we extracted morphological and colour descriptors (e.g., eccentricity, CD56 intensity), generating 19-dimensional nuclear and 29-dimensional gland feature vectors, with the latter also including epithelial organisation metrics.

Feature values were z-normalised before dimensionality reduction with UMAP. The resulting low-dimensional embeddings were partitioned into seven clusters for both glands and nuclei, defining distinct community memberships across the tissue landscape. At the patient level, targets comprised 13 temporally dependent gene expression measures and two derived expression ratios. Regression models were then applied independently to gland and nuclear compositions to quantify associations between structural cluster variation and molecular state.

#### Quantitative Feature-Gene Association Modelling

To assess associations between morphological features and gene expression, we implemented a regres-sion framework independent of communities. This model used the mean feature value aggregated across all segmented objects within each WSI to evaluate feature level changes linked to variations in gene expression. While numerous features achieved statistical significance, the majority exhibited modest effect sizes. Notably, features associated with the *GPX3/SLC15A2* ratio demonstrated the largest magnitudes (Figure S1). These results suggest the presence of a weak but biologically meaningful signal and reflect the intrinsic limitation of averaging across segmented objects, which diminishes information about spatial composition and object level heterogeneity.

#### Cluster Characterization and Interpretability Analysis

Following clustering, using our DCAFA framework, we investigated the defining characteristics of each cluster by modelling the binary cluster labels as instance level targets and performed regression analysis to identify enriched or depleted features. Across the six clusters retained for analysis (excluding the reference group), we observed distinct and biologically interpretable profiles. For example, among gland level clusters, Cluster 1 was characterised by densely packed epithelial cells positioned further from the gland boundary, and nuclear Cluster 6 displayed elevated CD56 expression, consistent with enrichment of uterine natural killer (uNK) cells (Figure 2). Representative examples of medoid proximal instances from each cluster illustrate the morphological profiles of each group and highlighting the interpretability of cluster assignments derived via DCAFA.

**Fig. 1:**
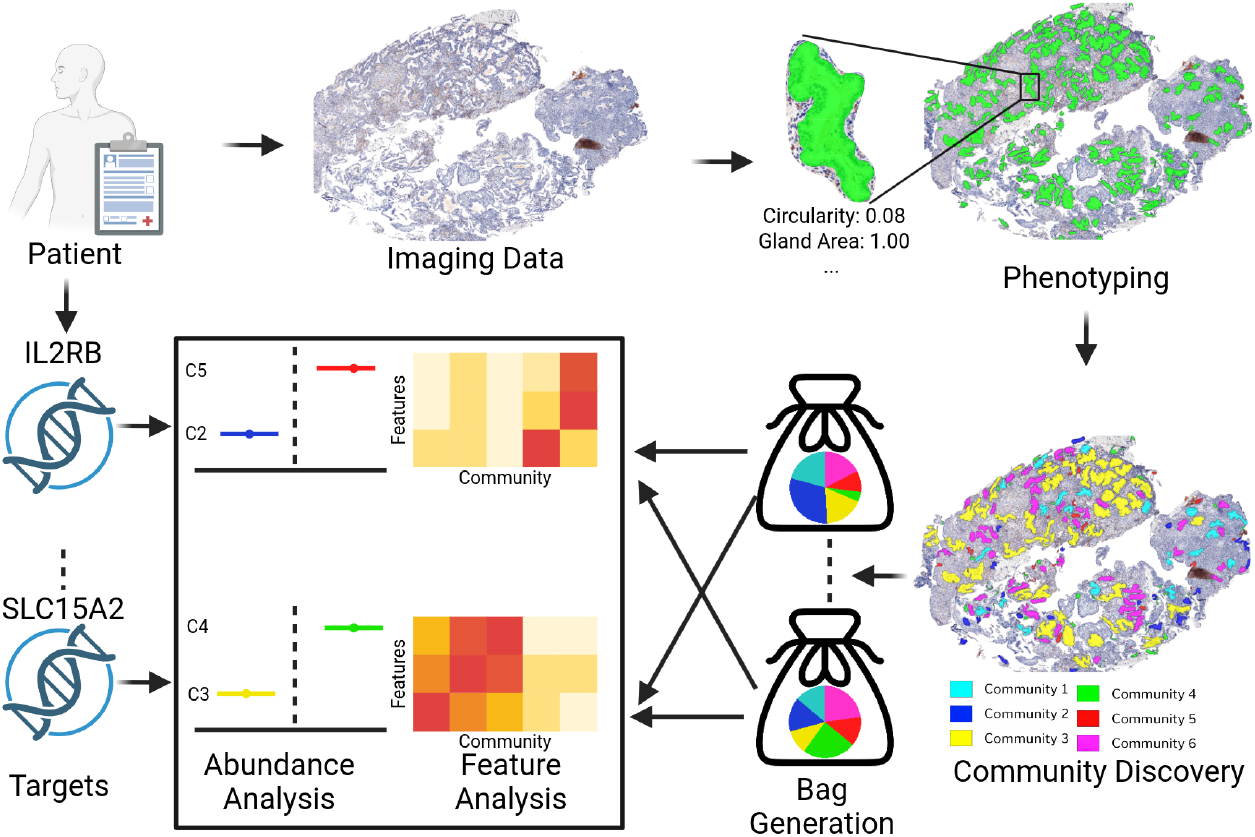
Overview of the DCAFA framework. For each patient, imaging data is collected and either manually annotated or processed using a pretrained phenotyping model (e.g., automated gland segmentation). Based on these segmentations, a community detection algorithm (e.g.k-means) is applied to assign instances to one or more communities. These instances are then represented as bags and integrated with patient-level or instance-level targets. This enables downstream analyses, including community abundance and feature analyses, to identify which features or communities are enriched or depleted relative to a target.

**Fig. 2:**
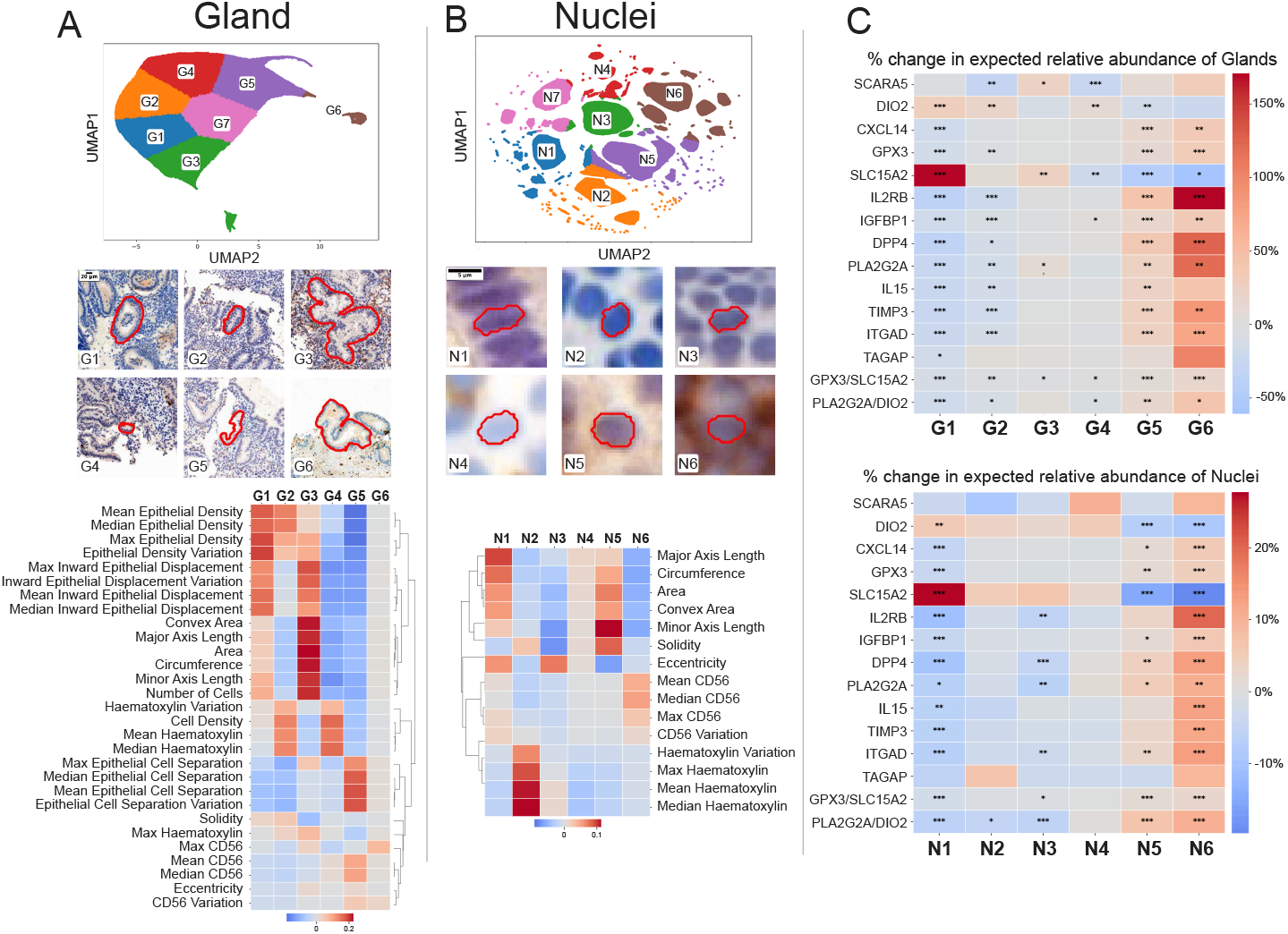
UMAP embeddings, representative cluster examples, and feature analyses presented as heatmaps showing regression coefficients derived from cluster labels against corresponding feature values, all demonstrating statistically significant patterns. Results are shown for (A) glandular and (B) nuclear communities. (C) Heatmaps display DCAFA exponentiated regression coefficients as a percentage increase indicating how each cluster varies with unit increase in gene expression levels for glands and nuclei. Positive coefficients (red) denote cluster enrichment, whereas negative coefficients (blue) indicate depletion with increasing target values. Statistical significance is denoted by: *:p *<* 0.05, **:p *<* 0.01, ***:p *<* 0.001.

#### Linking Tissue Community Composition to Gene Expression

To evaluate whether community abundances vary in relation to individual gene expression, we conducted a slide level differential community abundance analysis using the DCAFA framework. This model quantified associations between cluster proportions and slide-level expression of 13 genes within both glandular and nuclear compartments.

Glandular communities exhibited strong and biologically coherent relationships with canonical markers of endometrial maturation. Gland Cluster 1, correlated with high *SLC15A2* expression (exponentiated coefficient=2.707, 95% CI: 2.321–3.158), representing a 170% increase in abundance for each unit increase in gene expression. In contrast, Cluster 6 tracked high expression of *DPP4, GPX3*, and *CXCL14* (Figure 2C). Nuclear communities also revealed compartment specific expression signatures consistent with immune remodelling. The most striking pattern involved uNK cell differentiation markers *IL2RB* and *ITGAD*, which both exhibited high coefficients with nuclear Cluster 6, defined by their CD56 staining. We also show compositional changes with the ratios of genes associated with endometrial timing [12], and risk of miscarriage [8]. This demonstrates that DCAFA can uncover coordinated changes in endometrial tissue composition associated with gene expression, revealing patterns that are detectable from routine histopathological images.

### 3.2 Nuclear Morphological Correlates of Spatial Transcriptomic Signatures

Understanding cancer biology requires moving beyond individual genes to coordinated biological programmes, as many clinically relevant processes arise from the concerted activity of multiple genes. Analysing such pathways provides a more robust and interpretable view of tissue state, particularly in spatial settings where multiple processes coexist within local tumour regions. In spatial transcriptomics, these programmes can be identified in a data-driven manner as spatial transcriptomic signatures (STSs), which group genes that are consistently co-expressed within the same tissue patches. Each STS can therefore be interpreted as a latent biological pathway that is active or inactive across space.

However, while STSs capture molecular organisation, they do not directly indicate how these programmes manifest in histology or whether they correspond to clinically meaningful tissue phenotypes. Linking STSs to localised histological risk and cellular morphology is therefore essential, as it allows us to identify which transcriptional programmes are relevant to patient outcome and interpret them in terms of observable tissue structure.

To address this, we perform a three-stage analysis. First, we identify STSs associated with localised histological risk derived from the INSIGHT model, determining which programmes are most relevant to outcome. Second, we test whether variation in STS activity can be explained by differences in the abundance of morphologically defined nuclear communities. Finally, we perform instance-level feature analysis to characterise the morphological properties that define these communities, providing a direct interpretation of how transcriptional states manifest in histology [6].

We analysed the IMMUCAN cohort, consisting of 16 H&E-stained whole-slide images registered to a custom colorectal cancer spatial transcriptomic panel. Each tissue patch was annotated with 20 binary STS activation states (active/inactive) [6]. On the H&E images, we applied the Cerberus pipeline to segment approximately 600,000 nuclei [4], extracting eight interpretable morphological features describing nuclear size and shape.

#### Identifying Risk-Associated Spatial Transcriptomic Signatures

We first identified which STSs are associated with localised histological risk using patch-level outputs from the INSIGHT model. This analysis revealed STSs enriched in high- and low-risk regions. In Figure 3 i), STS8 (canonical differentiated epithelium), STS11 (regenerative epithelial differentiation), and STS13 (stem-like epithelial states) were strongly associated with higher-risk regions, consistent with their links to epithelial dedifferentiation and fetal-like transcriptional programmes [6].

**Fig. 3:**
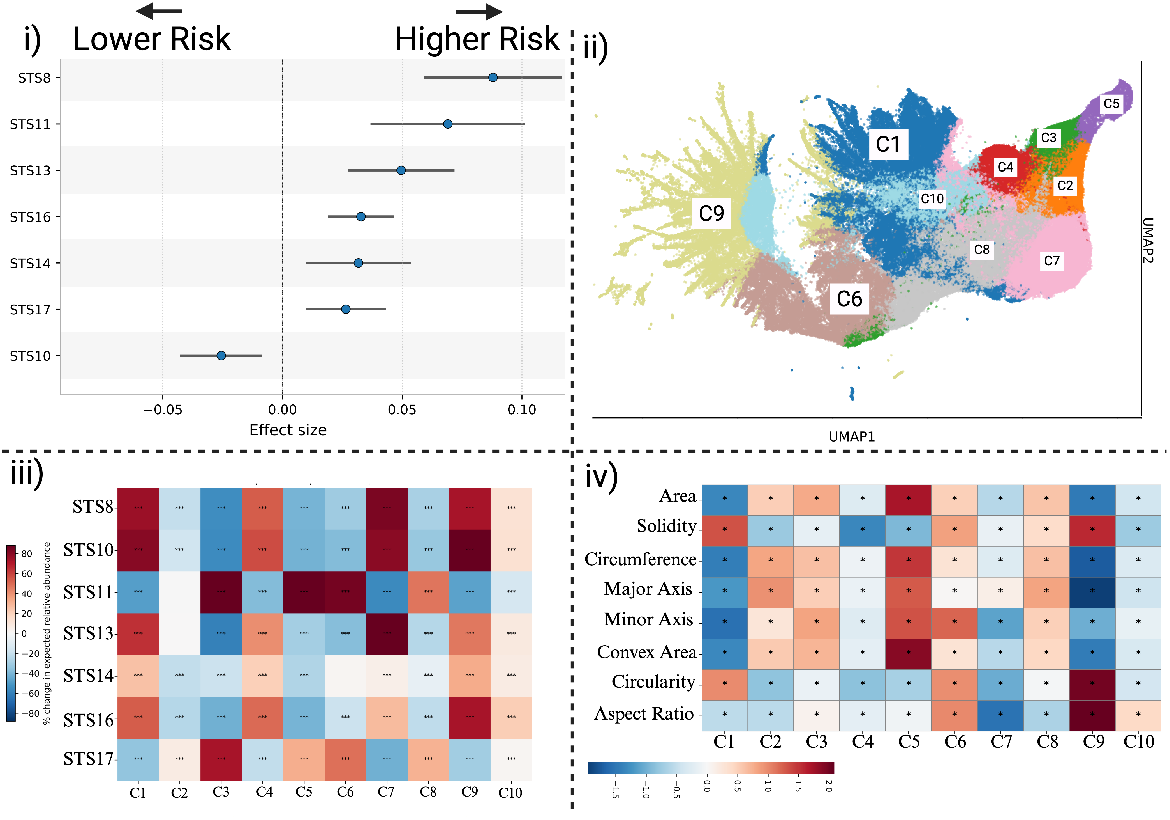
Spatial Transcriptomic Signature (STS) analysis workflow and results. (i) Association between individual STSs and histological risk, highlighting signatures significantly enriched in lower-versus higher-risk regions. (ii) UMAP visualization of nuclei clustered using k-means on automatically extracted cellular features. (iii) Heatmap showing the contribution of nuclei cluster abundances (C1–C10) to predicted patch-level STS values, indicating which cellular populations drive specific transcriptional signatures. (iv) Feature enrichment analysis across clusters, summarizing morphological characteristics that define each cluster and explain their association with STSs.

This step establishes which transcriptional programmes are clinically relevant and provides a principled basis for subsequent analysis.

#### Linking STS Activity to Nuclear Community Composition

We next investigated whether these transcriptional programmes could be explained by differences in tissue composition. Nuclei were clustered using *k*-means (*K* = 10) based on morphological features, defining nuclear communities representing distinct cell populations. Each patch (treated as a bag) was then represented by the abundance of these communities.

We modelled community abundance as a function of STS activation,adjusting for patient-level effects, testing whether specific programmes are associated with enrichment or depletion of particular nuclear populations. As shown in Figure 3 iii), we observed strong and structured associations between STSs and nuclear composition. For example, STS8 was enriched for clusters C1, C7, and C9, indicating that this programme corresponds to specific and recurring nuclear morphologies.

These results demonstrate that STSs reflect underlying shifts in the composition of histologically identifiable cell populations.

#### Morphological Characterisation of Nuclear Communities

To interpret these associations, we characterised the features defining each nuclear commu-nity using an instance-level generalised linear mixed model. This identified the morphological attributes most strongly associated with each cluster.

As shown in Figure 3 iv), clusters linked to high-risk STSs were characterised by distinct and interpretable traits. In particular, clusters C1 and C9 were defined by small, highly circular, and compact nuclei, consistent with features associated with high-risk regions. These morphological states may reflect chromatin stress linked to fetal-like transcriptional programmes [1].

Together, these results provide a coherent multi-scale interpretation of STSs, linking molecular programme activity to tissue composition and observable histological structure. This demonstrates how DCAFA enables mechanistic interpretation of spatial transcriptomic programmes directly from routine histology.

### 3.3 Additional Analyses

Additional results from other cohorts are provided in the Supplementary Material. In particular, these include: (i) an analysis of treatment response with respect to cell-type composition in H&E images from the Valentino cohort; (ii) an analysis of recurrence prediction using multiplex immunofluorescence expression in the Orion cohort; and (iii) an analysis of nuclear morphology in relation to point mutations in the TCGA-CRC cohort [11,7].

## 4 Discussion and Conclusion

Biological tissues comprise diverse cellular and structural elements whose spatial and compositional organisation underpins both physiological and pathological states. Translating these into quantitative, interpretable insights remains challenging, particularly when analytical methods overlook the inherent grouping of similar instances, such as morphologically related cells, glands, or tissue patches, whose relative abundances signal key biological processes.

Here, we introduced DCAFA, a unified statistical framework for analysing hierarchical biomedical image data by explicitly modelling composition as the primary analytical object. DCAFA bridges the gap between instance-level representations and higher-order tissue organisation, enabling both detection of compositionally distinct communities and identification of features that drive community membership and outcome associations.

Our formulation is flexible and applicable across diverse experimental designs. Empirical results across multiple domains including endometrial histopathology, colorectal cancer spatial transcriptomics, multiplex immunofluorescence, and predefined cell-type abundance analyses, demonstrate that DCAFA recovers biologically meaningful communities and provides interpretable, statistically sound characterisations of compositional and feature-level changes associated with disease states and treatment response.

DCAFA’s primary limitations include dependence on upstream clustering quality and computational demands for very high-dimensional feature spaces. Extreme compositional imbalance may also impact performance. Finally, all analyses are associative and do not imply causality. The regression formulation used is designed for stable inference but does not constitute a predictive model of outcomes, experimental validation is required for any discovered perturbations.

Collectively, this work establishes composition-aware modelling as a central paradigm in computational pathology and high-dimensional biomedical data analysis, and positions DCAFA as a general tool for translating complex, multi-level tissue representations into quantitative and clinically relevant insights.

